# Digital restoration of the pectoral girdles of two Early Cretaceous birds, and implications for early flight evolution

**DOI:** 10.1101/2021.12.10.472118

**Authors:** Shiying Wang, Yubo Ma, Qian Wu, Min Wang, Dongyu Hu, Corwin Sullivan, Xing Xu

## Abstract

Pectoral girdle morphology is a key determinant of flight capability, but in some respects is poorly known among stem birds. Here, we reconstruct the pectoral girdles of the Early Cretaceous birds *Sapeornis* and *Piscivorenantiornis* based on computed tomography and three-dimensional visualization, revealing key morphological details. Enantiornithines such as *Piscivorenantiornis* have a uniquely localized scapula-coracoid joint, with only one area of articulation. This single articulation contrasts with the double articulation widely present in non-enantiornithine pennaraptoran theropods, including *Sapeornis* and crown birds, which comprises main and subsidiary articular contacts. A partially closed triosseal canal occurs in non-euornithine birds, representing a transitional stage in flight apparatus evolution. Numerous modifications of the pectoral girdle along the line to crown birds, and lineage-specific pectoral girdle variations, produced diverse pectoral girdle morphologies among Mesozoic birds, which ensured that a commensurate range of capability levels and modes emerged during the early evolution of flight.

## Introduction

The evolution of powered flight in birds was one of the great transformations in vertebrate history, and involved a suite of dramatic anatomical changes that were needed to create a functional flight apparatus (*Dudley and Yanoviak, 2011; Padian, 1985; Rayner, 1988; Videler, 2005*). The pectoral girdle is a key component of the flight apparatus, and its function and evolutionary history have been extensively studied (*Baier et al., 2007; Bock, 2013; Novas et al., 2020; Senter, 2006*). However, most early bird fossils are essentially two-dimensionally preserved, and accordingly do not offer a full anatomical picture of the flight apparatus, a limitation that greatly hinders studies of early flight. Here, we reconstruct the pectoral girdles of the non-ornithothoracine bird *Sapeornis chaoyangensis* (PMoL-AB00015) and the enantiornithine bird *Piscivorenantiornis inusitatus* (IVPP V 22582) using computed tomography (CT) and three-dimensional visualization. Our reconstructions are the first three-dimensional ones of the pectoral girdle in any Cretaceous birds, and reveal some anatomical details that are important for understanding pectoral girdle evolution. For example, one main objective of our study was to better understand the evolution of the scapula-coracoid articulation and triosseal canal on the line to crown-group birds, because the form of the scapula-coracoid articulation is traditionally used to distinguish between enantiornithines and euornithines whereas the nature of the triosseal canal has implications for the course of the tendon of m. supracoracoideus and thus for the mechanics of the upstroke during flight.

## Results

Nearly complete shoulder girdles of *Sapeornis chaoyangensis* PMoL-AB00015 and *Piscivorenantiornis inusitatus* IVPP V 22582 have been reconstructed in detail (Figs. 1,2). However, the bones have been compressed during fossilization, so the reconstructions do not precisely capture the original morphology. Originally, for example, the furcula of *Sapeornis* PMoL-AB00015 was probably slightly curved caudally (despite being straight in our reconstruction), and the angle between the parts of the coracoid that contacted the scapula and the sternum was probably smaller than in our reconstruction. Because of these distortions, our reconstruction of *Sapeornis* is characterized by a larger distance between the two coracoids, and a more ventrally oriented glenoid fossa, than would have been present in the skeleton of the living animal. However, these effects do not affect our major conclusions.

**Figure 1.**
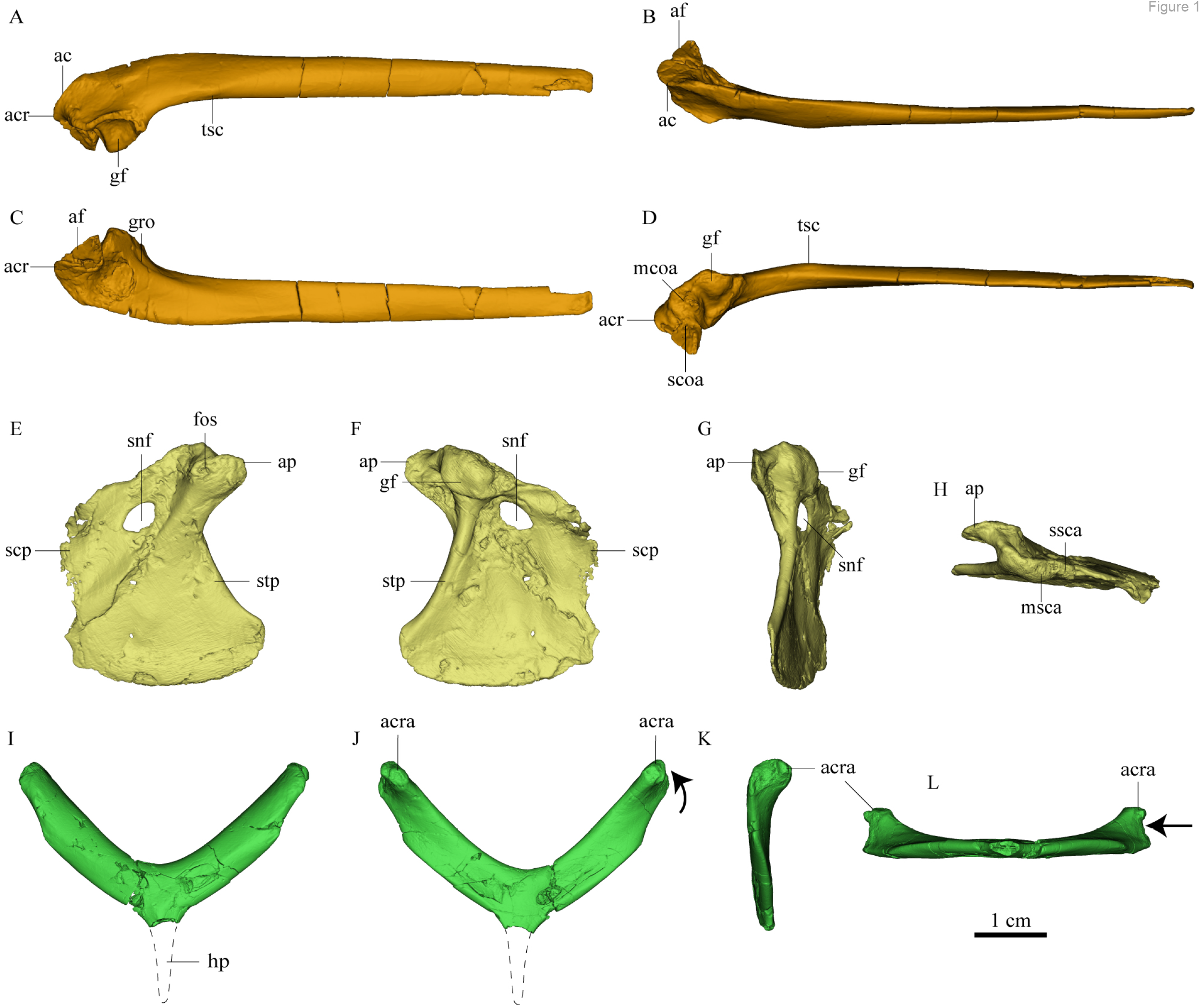
Pectoral girdle bones of *Sapeornis chaoyangensis* PMOL-AB00015. (**A-D**), left scapula in lateral, dorsal, medial (costal), and ventral views. (**E-H**), left coracoid in cranial, caudal, lateral, and proximal views; (**I-L**), furcula in cranial, caudal, lateral, and distal views. Abbreviations: ac, acromial crest; acr, acromion process; acra, articular surface for the acromion of the scapula; af, acromial flange; ap, acrocoracoid process; fos, fossa; gf, glenoid fossa; gro, groove of scapula; hp, hypocleidium; mcoa, main coracoidal articular surface; msca, main scapular articular surface; scoa, subsidiary coracoidal articular surface;scp, scapular part of coracoid; snf, supracoracoidal nerve foramen; ssca, subsidiary scapular articular surface; stp, sternal part of coracoid; tsc, tubercle of the scapula. The black arrows in J and L indicate the concave surface for the tendon of m. supracoracoideus.

**Figure 2.**
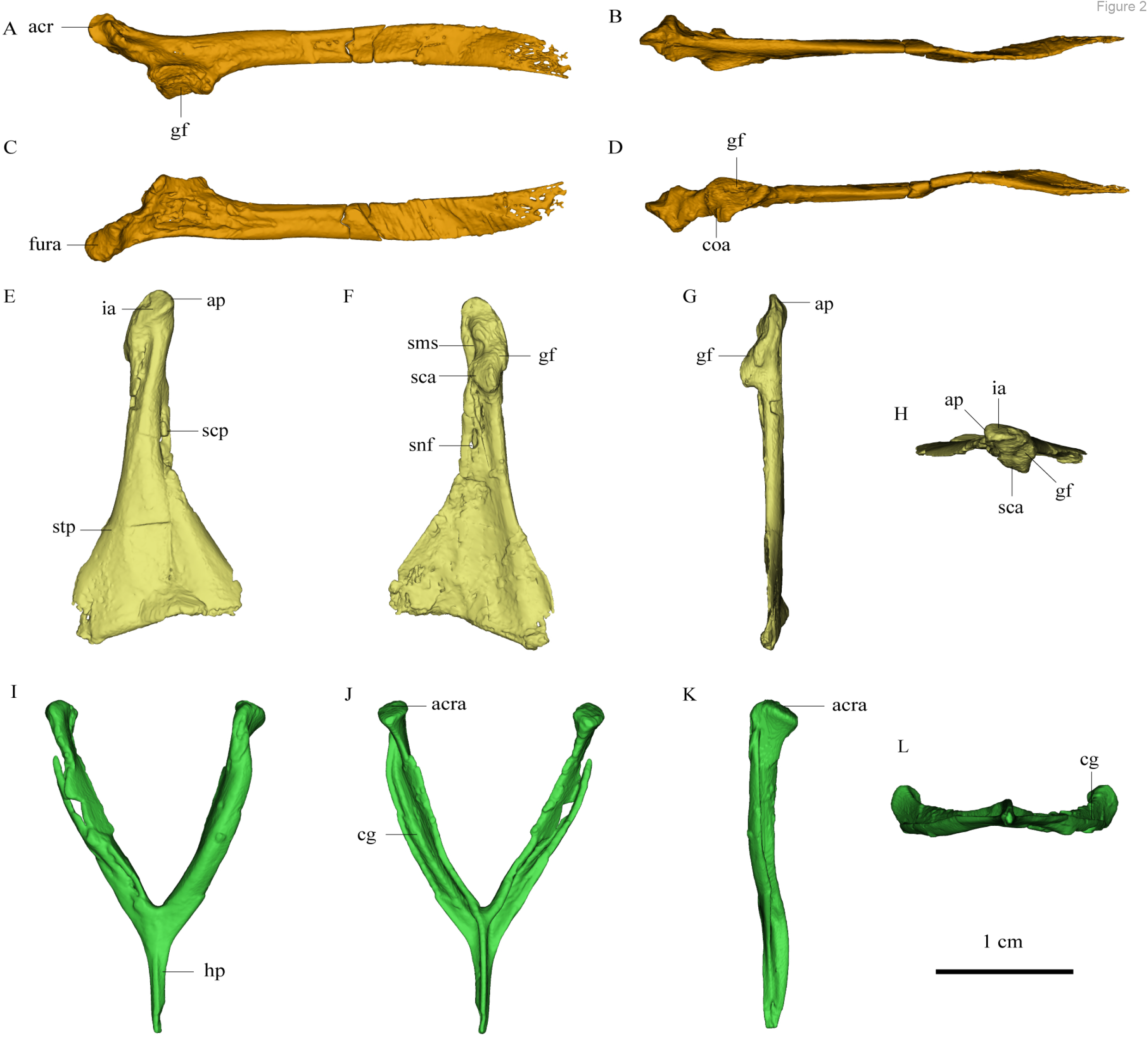
Pectoral girdle bones of *Piscivorenantiornis inusitatus* IVPP V 22582. (**A-D**), left scapula in lateral, dorsal, medial (costal), and ventral views. (**E-H**), right coracoid in cranial, caudal, lateral, and proximal views; (**I-L**), furcula in cranial, caudal, lateral, and distal views. Abbreviations: acr, acromion process; acra, articular surface for the acromion of the scapula; ap, acrocoracoid process; coa, coracoidal articular surface; cg, caudal groove; gf, glenoid fossa; hp, hypocleidium; ia, impression for the Lig. acrocoracohumerale; sca, scapular articular surface; scp, scapular part of coracoid; sms, sulcus m. supracoracoideus; snf, supracoracoidal nerve foramen; stp, sternal part of coracoid.

Osteology of the pectoral girdle of *Sapeornis* PMoL-AB00015. The proximal part of the left scapula is curved medially and ventrally. The scapular blade is slightly twisted about its longitudinal axis, to fit the curve of the ribcage as in extant birds (Fig. 1). The acromion is low and short. As in the dromaeosaurid *Rahonavis* (*Forster et al., 2020*), a broad flange protrudes costally from the acromion (Fig. 1, af). As in *Jeholornis* (*Zhou and Zhang, 2003*), the scapular glenoid fossa faces mainly ventrally but also slightly laterally, showing more lateral deflection than in deinonychosaurs (e.g., *Sinovenator* and *Rahonavis) (Forster et al., 2020)* and *Archaeopteryx (Zhou and Zhang, 2003*). The articular surface for the coracoid consists of two parts: a deeply concave main surface situated craniomedial to the glenoid facet (Fig. 3, msca) on the ventral surface of the scapula, and a more medially positioned subsidiary surface (Fig. 3, ssca). A weak tubercle lies on the ventrolateral margin of the scapula blade, possibly representing the muscle insertion site for the M. subscapulare (Fig. 1, tsc) (*Gianechini et al., 2018*). A short and shallow groove (Fig. 1, gro) on the medial surface of the scapula, close to the glenoid fossa and subparallel to the ventral margin, probably represents another muscle insertion site.

**Figure 3.**
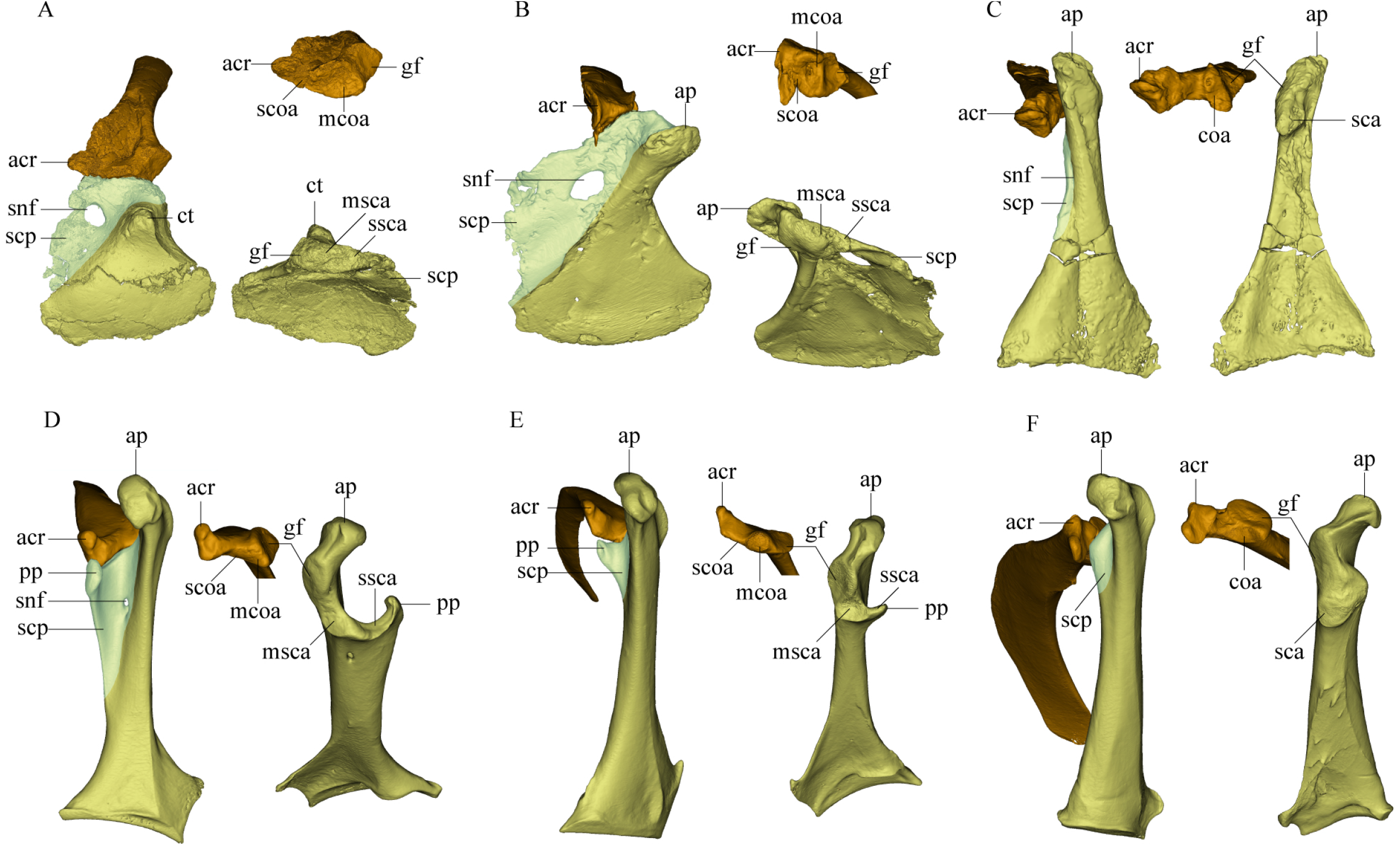
Comparison of scapula and coracoid morphology across various paravian taxa. Each panel shows articulated left scapula and coracoid in cranial view (on left) and opposing articular surfaces of left scapula and coracoid (on right, with cranial direction toward top of figure for both scapula and coracoid). (**A**), *Sinovenator changii* (mirrored); (**B**), *Sapeornis chaoyangensis*; (**C**), *Piscivorenantiornis inusitatus;* (**D**), *Tyto alba;* (**E**), *Egretta garzetta;* (**F**), *Pavo muticus.* Abbreviations: acr, acromion process; ap, acrocoracoid process; coa, coracoidal articular surface; ct, coracoid tubercle; gf, glenoid fossa; mcoa, main coracoidal articular surface; msca, main scapular articular surface; pp, procoracoid process; sca, scapular articular surface; scoa, subsidiary coracoidal articular surface; scp, scapular part of coracoid; snf, supracoracoidal nerve foramen; ssca, subsidiary scapular articular surface.

The coracoid is a broad, nearly rectangular plate. The distal margin of the coracoid is slightly convex and lacks an articular facet for the sternum (Fig. 1), rather than being straight to concave and bearing a sternal facet as in *Jeholornis (Wang et al., 2020)* and most ornithothoracines (*Atterholt et al., 2018; Wang and Zhou, 2017a*). The lack of a sternal facet lends further support to previous inferences that an ossified sternum is genuinely absent in this early pygostylian lineage (*Zheng et al., 2014*). On the cranial (ventral) surface of the coracoid, a ridge extends from the acrocoracoid process to the distomedial corner of the bone. This ridge divides the coracoid into separate portions that contact the sternum and the scapula (Fig. 1, stp and scp), as in deinonychosaurs (e.g., *Sinovenator*, *Mei*, and *Sinornithosaurus*) (*Xu, 2002*). The sternal part is short, the ratio of proximal-distal length to medial-lateral width at the distal margin being only 1.17. This is close to the value for *Archaeopteryx* (~1.15), but differs from those for the more elongated sternal rami of *Jeholornis*, *Confuciusornis,* and most ornithothoracines (generally > 1.5). The thin, sheet-like scapular part resembles the equivalent portion of the coracoid in some crown birds (e.g., *Tyto alba;* Fig. 3D), although this region gives rise to a narrow procoracoid process in most euornithines (Fig. 3E). The large oval supracoracoid foramen (Fig. 1, snf) is positioned within the scapular part of the coracoid and well distal to the scapular articular surface, as in non-avialan theropods.

The acrocoracoid process is short, blunt, and extends anteriorly only slightly beyond the level of the glenoid fossa as in *Jeholornis*, *Confuciusornis*, and enantiornithines (*Panteleev, 2018; Turner et al., 2012*). In most euornithines (e.g., Fig. 3E, D and F), by contrast, this process is proportionally longer, and extends much further beyond the glenoid, than in non-euornithine birds. In non-avialan theropods and *Archaeopteryx,* the coracoid tubercle (a homologue of the acrocoracoid process) is located anteroventral to the glenoid fossa (*Mayr et al., 2005; Novas et al., 2021*). The acrocoracoid process of *Sapeornis* forms a shelf-like structure projecting dorsally, cranially, and laterally from the lateralmost part of the coracoid, a condition not known in other birds. A small shallow fossa with an irregular surface, located at the medial end of the cranial surface of the acrocoracoid process (Fig. 1, fos), may have provided an attachment point for a coracoclavicular ligament connecting the coracoid and furcula (Fig. 4). In some volant extant birds, by contrast, the coracoclavicular ligament attaches to the dorsal edge of the medial surface of the acrocoracoid process (*Ghetie, 1976*).

**Figure 4.**
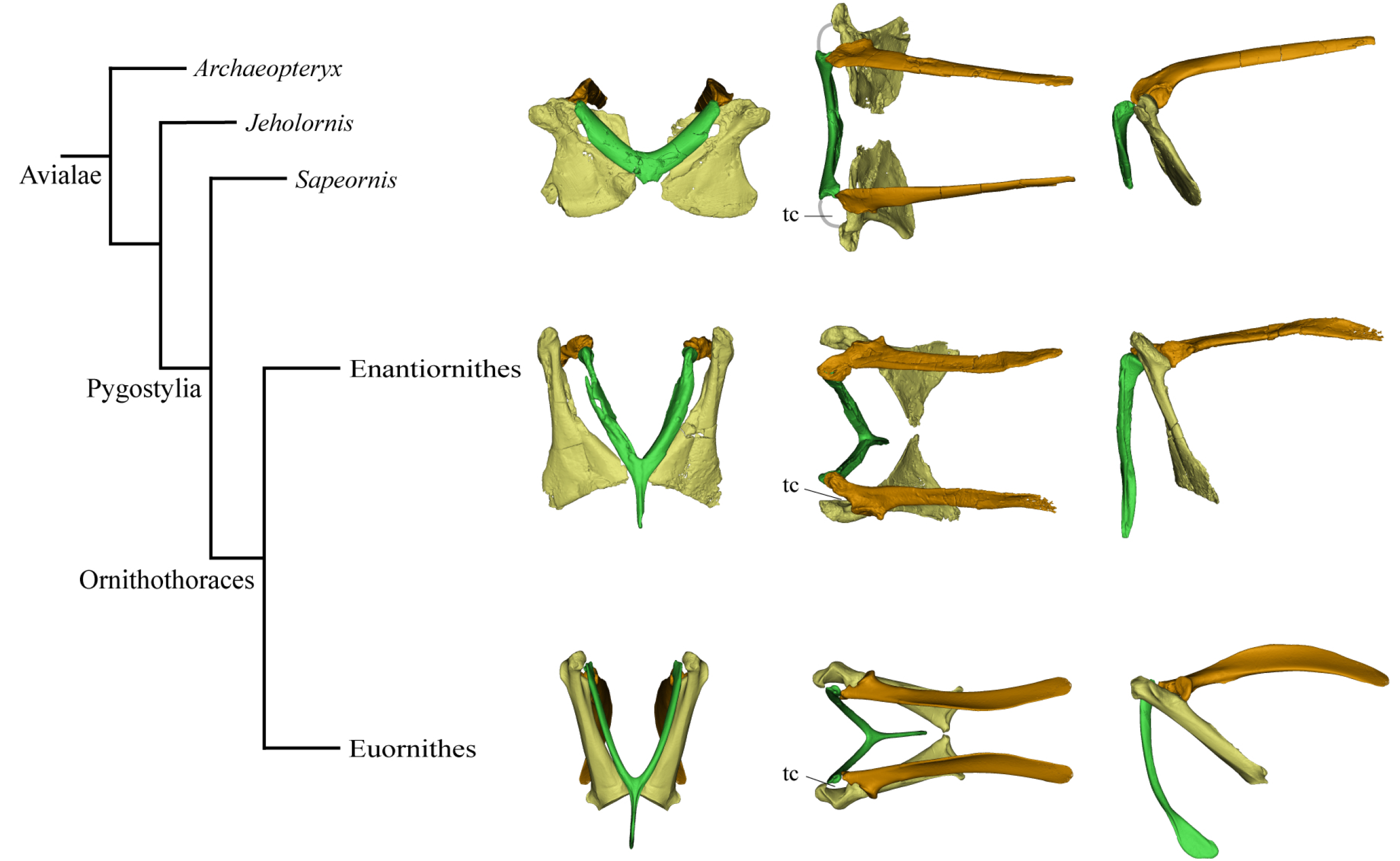
Simplified phylogeny with hypothetical steps of pectoral girdle evolution. The pectoral girdles of *Sapeornis chaoyangensis, Piscivorenantiornis inusitatus,* and *Pavo muticus* (from top to bottom) are shown in cranial, dorsal, and left lateral views. Abbreviations: tc, triosseal canal. The grey line in the dorsal view of the *Sapeornis* reconstruction represents the coracoclavicular ligament that connects the coracoid and furcula. Phylogenetic framework following Wang et al., (2018).

The glenoid fossa is on the proximolateral corner of the caudal face of the coracoid and wraps onto the proximal margin, whereas the scapular articular surface occupies the coracoid’s proximal margin, and there is no distinct border between the two articular areas. The coracoidal glenoid fossa is weakly concave and faces caudally and only slightly laterally, whereas the lateral tilt of the fossa is greater in late-diverging birds (Fig. 3).

Like the opposing surface on the scapula, the scapular articular surface on the coracoid is divided into main and subsidiary parts. The former is weakly convex, to match the concave main articular surface for the coracoid on the scapula, whereas the latter extends along the proximal margin of the scapular part of the coracoid and contacts the subsidiary coracoidal articular surface on the scapula.

The robust, craniocaudally compressed furcula could not have been as flexible as those of volant extant birds, which have mediolaterally compressed rami and in at least some cases act as a spring during flapping flight (*Boggs et al., 1997; Nesbitt et al., 2009*). The epicleidium has a blunt end and is slightly narrower than the ramus. In proximal view, the epicleidium is twisted by about 80° relative to the ramus. Similar torsion can also be observed in some other early birds (e.g., *Confuciusornis*), and in some deinonychosaurs (e.g., *Halszkaraptor, Buitreraptor) (Cau et al., 2021; Gianechini et al., 2018*) The lateral surface of the epicleidium is concave, making the cross-section of the omal end C-shaped. This concave surface possibly accommodated the tendon of the supracoracoideus muscle, which would have passed through the partially closed triosseal canal and over the small acrocoracoid process to reach its insertion on the humerus (Fig. 4). As in *Confuciusornis (Wu et al., 2020)* and *Xiaotingia (Xu et al., 2011),* a small caudal projection is located on the medial side of the omal tip, and probably articulated with the acromion of the scapula. The hypocleidium is broken away.

Osteology of the pectoral girdle of the *Piscivorenantiornis inusitatus* IVPP V 22582. The pectoral girdle is closely comparable in morphology to those of other enantiornithines. The long and robust acromion of the scapula is separated by a neck from the coracoidal articular surface. The cranial part of the medial surface of the acromion forms a flat articular surface for the furcula, as in many other enantiornithines (e.g., *Rapaxavis, Monoenantiornis,* and *Halimornis) (Chiappe et al., 2002; Chiappe et al., 2007; O’Connor et al., 201?*). The subtriangular scapular glenoid fossa faces more laterally than in *Sapeornis* and non-avialan theropods (*Forster et al., 2020),* although the orientation of the fossa is nevertheless also somewhat ventral. The slightly concave coracoidal articular surface lies cranial and costal to the scapular glenoid fossa, and is nearly perpendicular to the latter. The scapular blade is curved ventrally and tapered distally.

The coracoid is subtriangular in cranial or caudal view. The acrocoracoid process is minimally developed and rounded, as is typical in enantiornithines (*Panteleev, 2018*). The coracoidal glenoid fossa is proximodistally elongated and faces caudally and somewhat dorsolaterally. The slightly convex scapular articular surface is smaller than the coracoidal glenoid fossa, and is situated mediodistal to the latter as in other ornithothoracines. In *Sapeornis* and non-avialan theropods, the scapular articular surface is proportionally larger, and situated medial to the coracoidal glenoid fossa. On the medial side of the glenoid fossa, between the acrocoracoid process and scapular articular surface, lies a small incisure that is present in most enantiornithines (*Hu et al., 2015b; Panteleev, 2018; Wang et al., 2016d*). This structure is identified as the sulcus m. supracoracoideus, through which the m. supracoracoideus tendon passed (*Hu et al., 2012; Panteleev, 2018*). A large depression (Fig. 2, ia) associated with attachment of the Lig. acrocoracohumerale is located at the proximodistal level of the glenoid fossa, and faces cranially and slightly laterodorsally, whereas in extant birds the equivalent feature is located proximal to the glenoid fossa and faces laterally or laterodorsally. The remnant of the scapular part of the coracoid, termed the “medial crest” by Panteleev (2018), is extremely small and narrow, and terminates at the base of the scapular articular surface. It is perforated by a small supracoracoid foramen, as in most enantiornithines (*Atterholt et al., 2018; Chiappe et al., 2007; Panteleev, 2018; Wang et al., 2016a*). The shaft of the sternal part differs from the equivalent structure in most extant birds in being shorter, and in widening gradually over a considerable part of its length rather than abruptly near the sternal margin (*Panteleev, 2018*).

The furcula is robust and generally Y-shaped, with an interclavicular angle of only about 50° and a long hypocleidium, as in most enantiornithines (*Hu et al., 2015a; Zhang et al., 2014*). The omal tip is expanded both craniocaudally and mediolaterally to form a proximally facing articular facet for the coracoid, as in other enantiornithines (*Atterholt et al., 2018*). The midshaft of each ramus is ‘L’ shaped in cross-section owing to the presence of a deep caudal (dorsal) groove, another characteristic of Enantiornithes (*Atterholt et al., 2018; Chiappe et al., 2007; Close et al., 2009*). The proximal part of this groove faces somewhat laterally due to the torsion of the ramus, whereas the groove extends distally to the end of the hypocleidium, producing a high keel on the caudal surface of the hypocleidium between the right and left grooves. The bilaterally compressed form of the hypocleidium is shared with many enantiornithines, but differs from the craniocaudal compression seen in some oviraptorosaurs, some troodontids, and *Sapeornis* (*Nesbitt et al., 2009; Xu and Norell, 2004*).

## Discussion

*Sapeornis chaoyangensis* PMoL-AB00015 and *Piscivorenantiornis inusitatus* IVPP V 22582 provide significant new information on the pectoral girdle morphology of these two Early Cretaceous birds. This information bears, in particular, on the following issues pertaining to the early evolution of the avialan pectoral girdle.

### Morphology of the scapula-coracoid articulation, the scapular part of the coracoid, and the procoracoid process

The scapula-coracoid articulation is variable in morphology among pennaraptoran theropods. In non-avialan pennaraptorans, the structure of the articulation between the scapula and coracoid is not well known. This is mainly because the two elements tend to fuse, albeit normally at a relatively late ontogenetic stage, to form a scapulocoracoid. For example, the scapula and coracoid are tightly sutured together, but not fused, in a juvenile specimen of *Velociraptor* (MPC-D100/54), and a fused scapulocoracoid is seen in an adult specimen (IGM 100/986) (*Hone et al., 2012; Norell and Makovicky, 1999*). Complete fusion of the scapulocoracoid, leaving no visible suture, is the usual condition in adult individuals (e.g., *Citipati osmolskae* IGM 100/1004, *Anzu wyliei* CM 78 001) (*Lamanna et al., 2014; Norell et al., 2018*). Among non-ornithothoracine avialans, all known jinguofortisids, and all known confuciusornithiforms except a single subadult *Eoconfuciusornis zhengi* specimen (IVPP V 11977), possess a fused scapulocoracoid (*Wang et al., 2018; Wu et al., 2021*). The occurrence of scapula-coracoid fusion in *Archaeopteryx* is controversial, but the scapula and coracoid are separately preserved in some specimens (*Kundrát et al., 2019; Mayr et al., 2005; Wellnhofer et al., 2009; Wu et al., 2021*). Among ornithothoracines, a fused scapulocoracoid is known only in flightless paleognaths (*Wu et al., 2021*).

In several deinonychosaurs (e.g., *Sinovenator* and *Rahonavis),* the scapula and coracoid not only fail to fuse but remain loosely connected, contacting one another via smooth articular surfaces rather than via a tight, interdigitating suture as in other non-avialan pennaraptorans (*Forster et al., 2020*). In *Sinovenator* (Fig. 3A), the glenoid fossa of the coracoid is smaller than the scapular articular surface as in other non-avialan theropods. The reverse is true in crown group birds, and also in many stem birds in which this anatomical region is well known, such as *Piscivorenantiornis* (Fig. 2), *Elsornis (Chiappe et al., 2007), Mirarce (Atterholt et al., 2018)* and *Gansus (Wang et al., 2016c*). The scapular articular surface is located on the scapular part of the coracoid, and consists of two parts: a broadly convex main articular surface, and a flat to shallowly concave subsidiary articular surface. The corresponding coracoidal articular surfaces on the scapula are both concave. The main and subsidiary articulations together form a double articulation. *Bambiraptor* also has a double articulation, in which the smaller, shallowly convex subsidiary articular surface is located mediodorsally (medioproximally) to the flat main articular surface, and is separated from the latter by a low ridge. The corresponding coracoidal articular surfaces on the scapula are shallowly concave, as in *Sinovenator*. This implies a gap between the articular surfaces on the scapula and coracoid, which in life was presumably filled with cartilage. The unenlagiine *Rahonavis* possesses a double articulation with a concave main articular surface for the coracoid as in *Sinovenator* (*Forster et al., 2020*). As described above, the scapula-coracoid articulation of *Sapeornis* resembles that of *Sinovenator* in that the coracoid has a convex main articular surface for the scapula (matching the scapula’s concave main coracoidal articular surface) and a subsidiary articular surface that would have contacted the cranioventral margin of the acromion process of the scapula. *Sapeornis* thus has a double articulation, as in non-avialan pennaraptorans. In fact, avialans other than enantiornithines have a broadly uniform type of scapula-coracoid articulation, although some morphological variation is present. In *Jeholornis* and *Fukuipteryx* the coracoid has a concave main articular surface for the scapula (*Imai et al., 2019; Turner et al., 2012),* a feature that had been proposed as an apomorphy of the Euornithes (Ornithuromorpha) in previous studies (*Wang and Zhou, 2017a*). In most euornithines, the coracoid indeed has a deep cotyla that receives a corresponding convexity on the scapula (Fig. 3E). In some crown birds, however, the main scapular articular surface on the coracoid is flat to slightly convex (Fig. 3D and 3F), as in many enantiornithine birds and some non-avialan theropods such as *Sinovenator*. In most euornithines and *Jeholornis*, the subsidiary articular surface for the scapula is formed partly by the procoracoid process, which is a projection of the dorsomedial margin of the small scapular part of the coracoid.

Enantiornithines have a strikingly different scapula-coracoid articulation from other pennaraptorans. The presence of a single, convex articular surface fitting into the cotyla on the proximal end of the scapula has been widely accepted as a unique feature of the coracoid of enantiornithine birds (*Chiappe and Walker, 2002; Panteleev, 2018; Wang and Zhou, 2019*), because modern birds are considered to display the opposite condition, with the main scapular articular surface on the coracoid being concave. This purported discrepancy is the source of the clade name Enantiornithes, meaning “opposite birds”. As mentioned above, however, a convex scapular articular surface is also seen on the coracoids of some non-avialan theropods, such as *Sinovenator,* and even those of some crown birds (*Mayr, 2021)* though the coracoidal convexity is less prominent in these taxa than in late-diverging enantiornithine birds. Furthermore, the scapular articular surface on the coracoid is shallowly concave in some enantiornithines, such as pengornithids (e.g., IVPP V 18687 and V 18632). Consequently, the “opposite”-type scapula-coracoid articulation is neither present in all enantiornithine birds, nor unique to the Enantiornithes. However, our study indicates that the enantiornithine scapula-coracoid articulation is indeed unique, but for a different reason: absence or extreme reduction of the scapular part of the coracoid, and consequent loss of the subsidiary articular surface for the scapula in all enantiornithines (including pengornithids). This results in a single articulation in enantiornithines, in which the coracoid bears only one spatially restricted articular surface for the coracoid. The single articulation of enantiornithines is smaller in area, as well as morphologically simpler, than the double articulation of other pennaraptorans.

In summary, three important modifications to the scapula-coracoid articulation may be inferred to have occurred among pennaraptoran theropods: loss of fusion between the scapula and coracoid in the majority of adult avialans, a shift of the main articular surface for the scapula onto the sternal part of the coracoid in a clade comprising *Jeholornis* and pygostylians (reversed in *Sapeornis* to the primitive condition of having the main articular surface on the scapular part), and establishment of the unique single articulation by absence or extreme reduction of the scapular part of the coracoid in enantiornithine birds. All non-enantiornithine pennaraptorans, including crown birds, have a double articulation connecting the scapula and coracoid, and in the majority of non-enantiornithine birds, a procoracoid process is present to reinforce the double articulation and contribute to the triosseal canal. Previous assessments that enantiornithine and euornithine coracoids are neatly characterized by convex and concave articular surfaces for the scapula, respectively, are undermined by two observations. On the one hand, the main scapular articular surface on the coracoid is convex in some deinonychosaurs, non-ornithothoracine birds, and crown birds; on the other, a concave scapular cotyle on the coracoid is probably a plesiomorphic feature rather than an apomorphy of the Euornithes, as this condition is also seen in some non-ornithothoracine birds and pengornithid enantiornithines. The uniqueness of the enantiornithine coracoid does not lie primarily in the convexity of the articular surface for the scapula, but instead involves absence or extreme reduction of the scapular part of the bone, and consequent establishment of a single articulation.

### Architecture of the triosseal canal

The triosseal canal facilitates powered flapping flight in modern birds by forming a passage to admit the tendon of m. supracoracoideus, a muscle that contributes to humeral elevation and longitudinal rotation (*Baumel and Witmer, 1993; Poore et al., 1997*). The tendon ascends through the triosseal canal, passes laterally over the acrocoracoid process, which acts like a pulley to redirect the tendon, and ultimately inserts near the proximal end of the humerus. However, the name “triosseal canal” is misleading given the variable composition of this structure in living birds (*Livezey and Zusi, 2006*). The definition provided by Baumel and Witmer (1993) does not specify the architecture of the triosseal canal. In most extant birds the furcula, coracoid and scapula all participate (hence the name of the canal) in forming a fully enclosed bony passage. Typically, the epicleidium of the furcula forms the craniomedial wall of the triosseal canal, the acrocoracoid process of the coracoid forms the lateral wall, and the procoracoid process of the coracoid and the cranial margin of the acromion of the scapula forms the roof. However, in many species (e.g., *Phalacrocorax capillatus* and *Egretta garzetta*) the bony triosseal canal is only partially closed, in that the furcula lacks a bony contact with the scapula but is bound to the latter by Lig. scapuloclaviculare dorsale (*Baumel and Witmer, 1993*). In certain other birds a closed bony canal is formed by the coracoid and scapula only, with no contribution from the furcula, or even formed by the coracoid alone via an ossified bridge connecting the acrocoracoid and procoracoid processes (e.g., *Upupa epops* and *Columba livia*) (*Baumel and Witmer, 1993*). Therefore, the triosseal canal is not necessarily formed by all three pectoral elements, and not necessarily a fully enclosed bony passage. Also, the procoracoid process is absent in certain volant crown birds that possess a triosseal canal, including *Pavo muticus* (Fig. 4) and *Colius striatus* (*Mayr, 2021*), as well as in the Late Cretaceous galliform-like genus *Palintropus* (*Longrich, 2009*). Accordingly, the procoracoid process cannot be considered an essential constituent of the triosseal canal.

Among stem birds, the triosseal canal is widely accepted as present in early-diverging euornithines (*Mayr, 2017; Wang et al., 2016b; Zhou and Wang, 2017*). In most euornithine specimens, the acrocoracoid process is medially hooked and a prominent procoracoid process is present, features that suggest the existence of a typical, fully enclosed bony triosseal canal formed by the scapula, coracoid, and furcula. Many previous studies have denied the presence of a triosseal canal in non-euornithine birds because of the lack of a long medially hooked acrocoracoid process and a procoracoid process (*Novas et al., 2021; Wang and Zhou, 2017a*). Other studies have argued that a triosseal canal is present in enantiornithines, albeit based on limited evidence (*Kurochkin et al., 2013; Zhang and Zhou, 2000*). Mayr (2017) argued that the tendon of the m. supracoracoideus of enantiornithine birds ran along the medial side of the acromion, rather than along the lateral side as in crown birds, which would suggest that the supracoracoideus pulley system was differently configured in enantiornithines than in euornithines.

Our reconstructions of the pectoral girdles of *Sapeornis* and *Piscivorenantiornis* indicate the presence of a partially enclosed triosseal canal in these early stem birds. In *Sapeornis* the lateral wall of the triosseal canal is formed by the acrocoracoid process, the medial wall is formed by the furcula, and the roof is formed by the scapular part of the coracoid and the cranial margin of the acromion of the scapula. In these respects, the triosseal canal of *Sapeornis* has essentially the same structure as in most extant flying birds. In *Piscivorenantiornis,* the scapular part of the coracoid is extremely reduced and lacks a procoracoid process. The roof of the triosseal canal is formed by the cranioventral margin of the long acromion process of the scapula and the floor of the sulcus m. supracoracoideus of the coracoid, as in some extant birds that lack a procoracoid process, e.g., *Corvus corax, Pavo muticus* (Fig. 4). Both *Sapeornis* and enantiornithines have features of the pectoral girdle (e.g., dorsolaterally projecting acrocoracoid process in *Sapeornis*, and elongate acromion process and widely spaced tips of the coracoids in enantiornithines) implying the lack of a coracoid-furcula contact (*Mayr, 2017; Novas et al., 2021*). Absence of such a contact is a plesiomorphic feature, widely observed among non-avialan theropods (*Currie and Dong, 2001; Klingler, 2020; Lü, 2003*). Thus, *Sapeornis* and enantiornithines have only a partially enclosed bony triosseal canal, though a ligament could have completed the enclosure of the canal as in some modern birds (*Ghetie, 1976*).

### Pectoral girdle evolution and early flight

Mapping major aspects of pectoral girdle morphology onto an avialan phylogenetic tree suggests that three important evolutionary steps can be defined along the line to the modern flight apparatus, and reveals some distinctive features characterizing certain avialan clades (Fig. 4). Step I occurs at the base of the clade comprising *Jeholornis* and pygostylians, and involves torsion and elongation of scapular blade (ratio of scapular length to femoral length about 0.9, compared to 0.68 - 0.81 in *Archaeopteryx (Rauhut et al., 2018),* 0.68 in *Anchiornis (Hu et al., 2009)* and 0.58 - 0.73 in several dromaeosaurids (*Burnham et al., 2000; Hwang et al., 2002; Makovicky et al., 2005));* elongation of sternal part of coracoid (ratio of proximal-distal length of sternal part to medial-lateral width of distal margin of sternal part greater than 1.5; reversed to primitive condition in *Sapeornis);* shifting of scapular articular surface and glenoid fossa of coracoid to position extremely close to sternal part of bone; scapula-coracoid articulation localized, compared to condition in non-avialan theropods; extension of acrocoracoid process dorsal to level of coracoidal glenoid fossa; reduction in distance between left and right coracoids, indicating a relatively narrow and deep chest; reduction in angle between scapula and coracoid; and establishment of partially closed triosseal canal. Step II occurs at the base of Ornithothoraces, and involves cranial extension and thickening of acromion process; shifting of scapular articular surface and glenoid fossa of coracoid onto sternal part of bone; reduction in interclavicular angle (generally to less than 65°); and reduction of supracoracoidal nerve foramen. Step III occurs in early-diverging Euornithes, and involves downward curving of distal end of scapular blade; shifting of glenoid fossa of scapula onto external surface of bone, causing fossa to face dorsolaterally; appearance of procoracoid process on scapular part of coracoid; medial curving and further elongation of acrocoracoid process; further reduction in angle between scapula and coracoid; full enclosure of bony triosseal canal. Regarding distinctive pectoral girdle features in particular taxa, *Jeholornis* has an unusual combination of a prominent procoracoid process and a large supracoracoid foramen (*Lefèvre et al., 2014; Turner et al., 2012; Wang et al., 2020*), *Sapeornis* has a dorsolaterally oriented acrocoracoid process, and Enantiornithes is characterized by absence or at least further reduction of the scapular part of the coracoid, a single scapula-coracoid articulation, elongation of the hypocleidium, and presence of caudal grooves on the furcular rami and a keel on the caudal surface of the hypocleidium, and further solidification of furcula-scapula articulation.

The variation in pectoral girdle morphology seen among early birds is suggestive of a similarly wide diversity of flight capabilities and modes, an inference supported by previous studies (*Close and Rayfield, 2012; Heers and Dial, 2012; Novas et al., 2021*). The position of the acrocoracoid process, or coracoid tubercle, and the orientation of the glenoid fossa are functionally important, because the former is a key determinant of the course of the m. supracoracoideus tendon and the latter has a major effect of the range of motion of the wing (*Novas et al., 2020; Novas et al., 2021*). In *Archaeopteryx* and non-avialan pennaraptorans, the coracoid tubercle is located well below the level of the glenoid fossa and would not have acted as a pulley for the m. supracoracoideus tendon, so that m. supracoracoideus would have acted straightforwardly as a wing depressor (*Novas et al., 2020; Ostrom, 1974*). In the flightless paleognaths *Struthio* and *Rhea*, the acrocoracoid process is slightly below the coracoidal glenoid fossa and m. supracoracoideus is a protractor (*Novas et al., 2020*). The glenoid fossa of *Archaeopteryx,* non-avialan pennaraptorans, and flightless paleognaths faces laterally and has a sub-vertical major axis, indicating that the movements of the forelimb at the shoulder joint are, or in the case of extinct taxa would have been, predominantly cranial-caudal (*Novas et al., 2020; Ostrom, 1974*). In volant crown birds, by contrast, the well-developed acrocoracoid process is located further above the level of the glenoid fossa, and the fossa itself has a sub-horizontal major axis and faces laterodorsally (*Novas et al., 2020*). The m. supracoracoideus is the main elevator of the wing, and the wing moves more dorsoventrally at the shoulder (*Novas et al., 2020; Novas et al., 2021*).

In *Sapeornis* and most other non-ornithothoracine avialans, the acrocoracoid process is slightly higher than the coracoidal glenoid fossa, which would have caused the tendon of m. supracoracoideus to be slightly dorsally displaced relative to its position in *Archaeopteryx* and non-avialan pennaraptorans (*Mayr et al., 2005; Turner et al., 2012; Wang and Zhou, 2018*). The triosseal canal is located mediocranial to the glenoid fossa, whereas in extant birds the triosseal canal is located more directly medial to the glenoid fossa. In *Sapeornis* the vector of the tension exerted by m. supracoracoideus on the humerus is therefore directed craniodorsally, and promotes elevation, protraction, and pronation of the wing during the upstroke. The angle with the vertical formed by the major axis of the glenoid fossa is larger than in flightless paleognaths, but smaller than in volant extant birds (*Novas et al., 2020*). This suggests that the wing may have moved in a craniodorsal-caudoventral downstroke, unlike either the dorsal-ventral downstroke of extant volant birds or the largely cranial-caudal humeral movements of flightless paleognaths and presumably also of *Archaeopteryx* and non-avialan pennaraptorans. Several studies (*Mayr, 2017; Olson and Feduccia, 1979)* have suggested that the well-developed m. deltoideus which would have inserted broadly on the deltopectoral crest and humeral shaft, played the main role in the wing elevation in *Sapeornis* and other non-ornithothoracine birds. This seems consistent with the finding in this study that the m. supracoracoideus of *Sapeornis* would have pulled the humerus anteriorly (Fig. 4) rather than acting primarily as a wing elevator, and with the finding that Lig. acrocoracohumerale in *Sapeornis* had a relatively horizontal orientation, so that the dorsal shoulder musculature would have been largely responsible for preventing ventral dislocation of the humeral head when m. pectoralis was strongly activated. Nevertheless, the presence of the triosseal canal indicates that most non-ornithothoracine birds possessed some incipient capacity for powered, flapping flight. In *Piscivorenantiornis*, the acrocoracoid process is only slightly higher than the coracoidal glenoid fossa as in most non-ornithothoracine avialans, but nevertheless is considerably higher than the coracoid’s scapular articular surface as in euornithines, due to the proximodistally elongated coracoidal glenoid fossa. The orientation of the major axis of the glenoid fossa falls within the range seen in volant extant birds, and the triosseal canal is located medial to the glenoid fossa. Therefore, the wing movements of *Piscivorenantiornis* would have been more like those of volant extant birds than those of *Sapeornis,* indicating stronger flight capabilities in enantiornithines than in non-ornithothoracine birds.

In general, our study demonstrates some lineage-specific variations in pectoral girdle anatomy as well as significant modification of the pectoral girdle along the line to crown birds. The morphological diversity seen across the pectoral girdles of Mesozoic birds presumably resulted in a commensurate range of flight capabilities and modes in early flight evolution. The wing movements of *Sapeornis* would have differed from those of extant volant birds, highlighting the need to consider the possible effect of wing kinematics when reconstructing the flight ability of early birds.

## Materials and Methods

Institutional abbreviations: PMoL, Paleontological Museum of Liaoning, Shenyang, China; IVPP, Institute of Vertebrate Paleontology and Paleoanthropology, Chinese Academy of Sciences (CAS), Beijing, China.

*Sapeornis chaoyangensis* PMoL-AB00015 is a nearly complete semi-articulated skeleton collected from the Lower Cretaceous Jiufotang Formation at Yuanjiawa Village, Dapingfang Town, Chaoyang County, Liaoning Province, China. *Piscivorenantiornis inusitatus* IVPP V 22582 is a disarticulated skeleton collected from the Jiufotang Formation near Dapingfang Town (*Wang and Zhou, 2017b; Wang et al., 2016a*). These two specimens are the main data sources for this study. For comparison, we also examined the skeletons of several extant birds and fossil theropods, including *Tyto alba* IVPP OV 954, *Egretta garzetta* IVPP OV 1631, *Pavo muticus* IVPP OV 1668, the troodontid *Sinovenator changii* IVPP V 12615. The specimens were scanned using a GE v|tome|x m300&180 micro-computed-tomography scanner (GE Measurement & Control, Wuntsdorf, Germany) and a 225 kV micro-computerized tomography scanner (developed by the Institute of High Energy Physics, CAS), both housed at the Key Laboratory of Vertebrate Evolution and Human Origins of the Chinese Academy of Sciences. 3D segmentation of the CT data was performed using the software package Mimics (19.0).

## Acknowledgments

We acknowledge Xiaoqing Ding for specimen preparation; Yun Feng and Yemao Hou for help with CT scanning; Paul Rummy for discussion.

## References

Atterholt J, Hutchison JH, and O’Connor JK. 2018. The most complete enantiornithine from North America and a phylogenetic analysis of the Avisauridae. PeerJ 6:e5910. doi:10.7717/peerj.5910.

Baier DB, Gatesy SM, and Jenkins FA. 2007. A critical ligamentous mechanism in the evolution of avian flight. Nature 445:307–310. doi:10.1038/nature05435.

Baumel JJ, and Witmer LM. 1993. Osteologia. In: Baumel JJ, King AS, Breazile JE, Evans HE, and Vanden Berge JC, editors. Handbook of avian anatomy: nomina anatomica avium 2nd ed: Publications of the Nuttall Ornithological Club. p 45–124.

Bock WJ. 2013. The furcula and the evolution of avian flight. Paleontological Journal 47:1236–1244. doi:10.1134/s0031030113110038.

Boggs DF, Farish A Jenkins J, and Dia KP. 1997. The effects of the wingbeat cycle on respiration in black-billed magpies (*Pica pica*). Journal Of Experimental Biology 200:1403–1412.

Burnham DA, Derstler KL, Currie PJ, Bakker RT, and Ostrom JH. 2000. Remarkable new birdlike dinosaur (Theropoda: Maniraptora) from the Upper Cretaceous of Montana. University of Kansas Paleontological Contributions 13:1–14.

Cau A, Beyrand V, Barsbold R, Tsogtbaatar K, and Godefroit P. 2021. Unusual pectoral apparatus in a predatory dinosaur resolves avian wishbone homology. Sci Rep 11:14722. doi:10.1038/s41598-021-94285-3.

Chiappe LM, Lamb JP, Jr., and Ericson PGP. 2002. New enantiornithine bird from the marine Upper Cretaceous of Alabama. Journal of Vertebrate Paleontology 22:170–174.

Chiappe LM, Suzuki S, Dyke GJ, Watabe M, Tsogtbaatar K, and Barsbold R. 2007. A new enantiornithine bird from the Late Cretaceous of the Gobi desert. Journal of Systematic Palaeontology 5:193–208. doi:10.1017/s1477201906001969.

Chiappe LM, and Walker CA. 2002. Skeletal morphology and systematics of the Cretaceous Euenantiornithes (Ornithothoraces: Enantiornithes). In: Chiappe LM, and Witmer LM, editors. Mesozoic birds: above the heads of dinosaurs. Berkeley, California: University of California Press. p 559–588.

Close RA, and Rayfield EJ. 2012. Functional morphometric analysis of the furcula in Mesozoic birds. PLoS One 7:e36664. doi:10.1371/journal.pone.0036664.

Close RA, Vickers-Rich P, Trusler P, Chiappe LM, O’Connor J, Rich TH, Kool L, and Komarower P. 2009. Earliest Gondwanan bird from the Cretaceous of southeastern Australia. Journal of Vertebrate Paleontology 29:616–619.

Currie PJ, and Dong Z. 2001. New information on Cretaceous troodontids (Dinosauria, Theropoda) from the People’s Republic of China. Canadian Journal of Earth Sciences 38:1753–1766. doi:10.1139/e01-065.

Dudley R, and Yanoviak SP. 2011. Animal aloft: the origins of aerial behavior and flight. Integrative and Comparative Biology 51:926–936. doi:10.1093/icb/icr002.

Forster C, O’Connor P, Chiappe L, and Turner A. 2020. The osteology of the Late Cretaceous paravian *Rahonavis ostromi* from Madagascar. Palaeontologia Electronica 23:a31. doi:10.26879/793.

Ghetie V. 1976. Anatomical atlas of domestic birds. Bucharest: Editura Academiei Republich Socialiste Romania. p80.

Gianechini FA, Makovicky PJ, Apesteguia S, and Cerda I. 2018. Postcranial skeletal anatomy of the holotype and referred specimens of *Buitreraptor gonzalezorum* Makovicky, Apesteguía and Agnolín 2005 (Theropoda, Dromaeosauridae), from the Late Cretaceous of Patagonia. PeerJ 6:e4558. doi:10.7717/peerj.4558.

Heers AM, and Dial KP. 2012. From extant to extinct: locomotor ontogeny and the evolution of avian flight. Trends Ecol Evol 27:296–305. doi:10.1016/j.tree.2011.12.003.

Hone D, Tsuihiji T, Watabe M, and Tsogtbaatr K. 2012. Pterosaurs as a food source for small dromaeosaurs. Palaeogeography, Palaeoclimatology, Palaeoecology 331-332:27–30. doi:10.1016/j.palaeo.2012.02.021.

Hu D, Hou L, Zhang L, and Xu X. 2009. A pre-*Archaeopteryx* troodontid theropod from China with long feathers on the metatarsus. Nature 461:640–643. doi:10.1038/nature08322.

Hu D, Liu Y, Li J, Xu X, and Hou L. 2015a. *Yuanjiawaornis viriosus*, gen. et sp. nov., a large enantiornithine bird from the Lower Cretaceous of western Liaoning, China. Cretaceous Research 55:210–219. doi:10.1016/j.cretres.2015.02.013.

Hu D, Xu X, Hou L, and Sullivan C. 2012. A new enantiornithine bird from the Lower Cretaceous of Western Liaoning, China, and its implications for early avian evolution. Journal of Vertebrate Paleontology 32:639–645. doi:10.1080/02724634.2012.652321.

Hu H, O’Connor JK, and Zhou Z. 2015b. A new species of Pengornithidae (Aves: Enantiornithes) from the Lower Cretaceous of China suggests a specialized scansorial habitat previously unknown in early birds. PLoS One 10:e0126791. doi:10.1371/journal.pone.0126791.

Hwang SH, Norell M, Ji Q, and Gao K. 2002. New specimens of *Microraptor zhaoianus* (Theropoda, Dromaeosauridae) from northeastern China. American Museum Novitates 3381:1–44.

Imai T, Azuma Y, Kawabe S, Shibata M, Miyata K, Wang M, and Zhou Z. 2019. An unusual bird (Theropoda, Avialae) from the Early Cretaceous of Japan suggests complex evolutionary history of basal birds. Communicating Biology 2:399. doi:10.1038/s42003-019-0639-4.

Klingler JJ. 2020. The evolution of the pectoral extrinsic appendicular and infrahyoid musculature in theropods and its functional and behavioral importance. Journal of Anatomy 237:870–889. doi:10.1111/joa.13256.

Kundrát M, Nudds J, Kear BP, Lü J, and Ahlberg P. 2019. The first specimen of *Archaeopteryx* from the Upper Jurassic Mörnsheim Formation of Germany. Historical Biology 31:3–63. doi:10.1080/08912963.2018.1518443.

Kurochkin EN, Chatterjee S, and Mikhailov KE. 2013. An embryonic enantiornithine bird and associated eggs from the cretaceous of Mongolia. Paleontological Journal 47:1252–1269. doi:10.1134/s0031030113110087.

Lamanna MC, Sues HD, Schachner ER, and Lyson TR. 2014. A new large-bodied oviraptorosaurian theropod dinosaur from the latest Cretaceous of western North America. PLoS One 9:e92022. doi:10.1371/journal.pone.0092022.

Lefèvre U, Hu D, Escuillié F, Dyke G, and Pascal G. 2014. A new long-tailed basal bird from the Lower Cretaceous of north-eastern China. Biological Journal of the Linnean Society. p790–804.

Livezey BC, and Zusi RL. 2006. Phylogeny of Neornithes. Bulletin of Carnegie Museum of Natural History 37:1–544. doi:10.2992/0145-9058(2006)37[1:Pon]2.0.Co;2.

Longrich N. 2009. An ornithurine-dominated avifauna from the Belly River Group (Campanian, Upper Cretaceous) of Alberta, Canada. Cretaceous Research 30:161–177. doi:10.1016/j.cretres.2008.06.007.

Lü J. 2003. A new oviraptorosaurid (Theropoda: Oviraptorosauria) from the Late Cretaceous of southern China. Journal of Vertebrate Paleontology 22:871–875. doi:10.1671/0272-4634(2002)022[0871:Anotof]2.0.Co;2.

Makovicky PJ, Apesteguia S, and Agnolin FL. 2005. The earliest dromaeosaurid theropod from South America. Nature 437:1007–1011. doi:10.1038/nature03996.

Mayr G. 2017. Pectoral girdle morphology of Mesozoic birds and the evolution of the avian supracoracoideus muscle. Journal of Ornithology 158:859–867. doi:10.1007/s10336-017-1451-x.

Mayr G. 2021. The coracoscapular joint of neornithine birds—extensive homoplasy in a widely neglected articular surface of the avian pectoral girdle and its possible functional correlates. Zoomorphology 140:217–228. doi:org/10.1007/s00435-021-00528-2.

Mayr G, Pohl B, and Peters DS. 2005. A well-preserved *Archaeopteryx* specimen with theropod features. Science 310:1483–1486. doi:10.1126/science.1120331.

Nesbitt SJ, Turner AH, Spaulding M, Conrad JL, and Norell MA. 2009. The theropod furcula. Journal Of Morphology 270:856–879. doi:10.1002/jmor.10724.

Norell MA, Balanoff AM, Barta DE, and Erickson GM. 2018. A second specimen of Citipati osmolskae associated with a nest of eggs from Ukhaa Tolgod, Omnogov Aimag, Mongolia. 3899:1–45. doi:10.5962/bhl.title.156722.

Norell MA, and Makovicky PJ. 1999. Important features of the dromaeosaurid skeleton II: Information from newly collected specimens of *Velociraptor mongoliensis*. American Museum Novitates 3282:1–45.

Novas FE, Agnolin FL, Egli FB, and Lo Coco GE. 2020. Pectoral girdle morphology in early-diverging paravians and living ratites: implications for the origin of flight. In: Pittman M, and Xu X, editors. Pennaraptoran theropod dinosaurs past progress and new frontiers: Bulletin of the American Museum of Natural History. p345–353.

Novas FE, Motta MJ, Agnolin FL, Rozadilla S, Lo Coco GE, and Egli FB. 2021. Comments on the morphology of basal paravian shoulder girdle: new data based on unenlagiid theropods and paleognath birds. Frontiers in Earth Science 9:1–14. doi:ARTN 66216710.3389/feart.2021.662167.

O’Connor JK, Chiappe LM, Gao C, and Zhao B. 2011. Anatomy of the Early Cretaceous enantiornithine bird *Rapaxavis pani*. Acta Palaeontologica Polonica 56:463–475. doi:10.4202/app.2010.0047.

Olson SL, and Feduccia A. 1979. Flight capability and the pectoral girdle of *Archaeopteryx*. Nature 278:247–248.

Ostrom JH. 1974. *Archaeopteryx* and the origin of flight. Quarterly Review of Biology 49:27–47.

Padian K. 1985. The origins and aerodynamics of flight in extinct vertebrates. Palaeontology 28:413–433.

Panteleev AV. 2018. Morphology of the coracoid of Late Cretaceous enantiornithines (Aves: Enantiornithes) from Dzharakuduk (Uzbekistan). Paleontological Journal 52:201–207. doi:10.1134/s0031030118020089.

Poore SO, Sanchez-Haiman A, and Goslow Jr GE. 1997. Wing upstroke and the evolution of flapping flight. Nature 387:799–802. doi:10.1038/42930.

Rauhut OWM, Foth C, and Tischlinger H. 2018. The oldest *Archaeopteryx* (Theropoda: Avialiae): a new specimen from the Kimmeridgian/Tithonian boundary of Schamhaupten, Bavaria. PeerJ 6:e4191. doi:10.7717/peerj.4191.

Rayner JMV. 1988. The evolution of vertebrate flight. Biological Journal of the Linnean Society 34:269–287.

Senter P. 2006. Scapular orientation in theropods and basal birds and the origin of flapping flight. Acta Palaeontologica Polonica 51:305–313.

Turner AH, Makovicky PJ, and Norell MA. 2012. A review of dromaeosaurid systematics and paravian phylogeny. Bulletin of the American Museum of Natural History 371:1–206.

Videler JJ. 2005. Avian Flight. New York: Oxford University Press. p26–231.

Wang M, Stidham TA, and Zhou Z. 2018. A new clade of basal Early Cretaceous pygostylian birds and developmental plasticity of the avian shoulder girdle. Proc Natl Acad Sci U S A 115:10708–10713. doi:10.1073/pnas.1812176115.

Wang M, and Zhou Z. 2017a. The evolution of birds with implications from new fossil evidences. In: Maina JN, editor. The biology of the avian respiratory system. Cham: United Kingdom: Springer Nature. p 1–26.

Wang M, and Zhou Z. 2017b. A morphological study of the first known piscivorous enantiornithine bird from the Early Cretaceous of China. Journal of Vertebrate Paleontology 37:e1278702. doi:10.1080/02724634.2017.1278702.

Wang M, and Zhou Z. 2018. A new confuciusornithid (Aves: Pygostylia) from the Early Cretaceous increases the morphological disparity of the Confuciusornithidae. Zoological Journal of the Linnean Society 185:417–430. doi:10.1093/zoolinnean/zly045/5066665.

Wang M, and Zhou Z. 2019. A new enantiornithine (Aves: Ornithothoraces) with completely fused premaxillae from the Early Cretaceous of China. Journal of Systematic Palaeontology 17:1299–1312. doi:10.1080/14772019.2018.1527403.

Wang M, Zhou Z, and Sullivan C. 2016a. A fish-eating enantiornithine bird from the Early Cretaceous of China provides evidence of modern avian digestive features. Curr Biol 26:1170–1176. doi:10.1016/j.cub.2016.02.055.

Wang M, Zhou Z, and Zhou S. 2016b. A new basal ornithuromorph bird (Aves: Ornithothoraces) from the Early Cretaceous of China with implication for morphology of early Ornithuromorpha. Zoological Journal of the Linnean Society 176:207–223. doi:10.1111/zoj.12302.

Wang X, Huang J, Kundrát M, Cau A, Liu X, Wang Y, and Ju S. 2020. A new jeholornithiform exhibits the earliest appearance of the fused sternum and pelvis in the evolution of avialan dinosaurs. Journal of Asian Earth Sciences 199:1–18. doi:10.1016/j.jseaes.2020.104401.

Wang Y-M, O’Connor JK, Li D-Q, and You H-L. 2016c. New information on postcranial skeleton of the Early Cretaceous *Gansus yumenensis* (Aves: Ornithuromorpha). Historical Biology 28:666–679. doi:10.1080/08912963.2015.1006217.

Wang Y, Wang M, O’Connor JK, Wang X, Zheng X, and Zhang X. 2016d. A new Jehol enantiornithine bird with three-dimensional preservation and ovarian follicles. Journal of Vertebrate Paleontology 36:e1054496. doi:10.1080/02724634.2015.1054496.

Wellnhofer P, Haase F, and Chiappe LM. 2009. Archaeopteryx: the icon of evolution. Munich: Verlag Dr. Friedrich Pfeil. p57–172.

Wu Q, Bailleul AM, Li Z, O’Connor J, and Zhou Z. 2021. Osteohistology of the scapulocoracoid of *Confuciusornis* and preliminary analysis of the shoulder Joint in Aves. Frontiers in Earth Science 9:617124. doi:10.3389/feart.2021.617124.

Wu Q, O’Connor J, Li Z, and Bailleul AM. 2020. Cartilage on the furculae of living birds and the extinct bird *Confuciusornis*: a preliminary analysis and implications for flight style inferences in Mesozoic birds. Vertebrata PalAsiatica 59:106–124. doi:10.19615/j.cnki.1000-3118.201222.

Xu X. 2002. Deinonychosaurian fossils from the Jehol Group of western Liaoning and the coelurosaurian evolution Ph.D. Dissertation. Beijing: Chinese Academy of Sciences.

Xu X, and Norell M. 2004. A new troodontid dinosaur from China with avian-like sleeping posture. Nature 431:838–841. doi:10.1038/nature02898.

Xu X, You H, Du K, and Han F. 2011. An *Archaeopteryx*-like theropod from China and the origin of Avialae. Nature 475:465–470. doi:10.1038/nature10288.

Zhang F, and Zhou Z. 2000. A primitive enantiornithine bird and the origin of feathers. Science 290:1955–1959. doi:10.1126/science.290.5498.1955.

Zhang Y, O’Connor J, Liu D, Meng Q, Sigurdsen T, and Chiappe LM. 2014. New information on the anatomy of the Chinese Early Cretaceous Bohaiornithidae (Aves: Enantiornithes) from a subadult specimen of *Zhouornis hani*. PeerJ 2:e407. doi:10.7717/peerj.407.

Zheng X, O’Connor J, Wang X, Wang M, Zhang X, and Zhou Z. 2014. On the absence of sternal elements in *Anchiornis* (Paraves) and *Sapeornis* (Aves) and the complex early evolution of the avian sternum. Proc Natl Acad Sci U S A 111:13900–13905. doi:10.1073/pnas.1411070111.

Zhou Z-H, and Wang Y. 2017. Vertebrate assemblages of the Jurassic Yanliao Biota and the Early Cretaceous Jehol Biota: comparisons and implications. Palaeoworld 26:241–252. doi:10.1016/j.palwor.2017.01.002.

Zhou Z, and Zhang F. 2003. *Jeholornis* compared to *Archaeopteryx*, with a new understanding of the earliest avian evolution. Naturwissenschaften 90:220–225. doi:10.1007/s00114-003-0416-5.

